# High-resolution regulatory maps connect cardiovascular risk variants to disease related pathways

**DOI:** 10.1101/376699

**Authors:** Örjan Åkerborg, Rapolas Spalinskas, Sailendra Pradhananga, Anandashankar Anil, Pontus Höjer, Flore-Anne Poujade, Lasse Folkersen, Per Eriksson, Pelin Sahlén

**Affiliations:** Science for Life Laboratory, School of Engineering Sciences in Chemistry, Biotechnology and Health, Division of Gene Technology, KTH Royal Institute of Technology, Solna, Sweden; Cardiovascular Medicine Unit, Center for Molecular Medicine, Department of Medicine, Karolinska Institutet, Stockholm, Sweden; Department of Bioinformatics, Technical University of Denmark, Copenhagen, Denmark

**Author notes:** Share first authorship (Örjan Åkerborg, Rapolas Spalinskas, Sailendra Pradhananga).

**Keywords:** Gene regulation, Promoter-enhancer interaction, HiCap, GWAS, variants, Linkage disequilibrium

## Abstract

Genetic variant landscape of cardiovascular disease (CVD) is dominated by non-coding variants among which many occur within putative enhancers regulating the expression levels of relevant genes. It is crucial to assign the genetic variants to their correct gene both to gain insights into perturbed functions and better assess the risk of disease. In this study, we generated high-resolution genomic interaction maps (~750 bases) in aortic endothelial, smooth muscle and THP-1 macrophages using Hi-C coupled with sequence capture targeting 25,429 features including variants associated with CVD. We detected interactions for 761 CVD risk variants obtained by genome-wide association studies (GWAS) and identified novel as well as established functions associated with CVD. We were able to fine-map 331 GWAS variants using interaction networks, thereby identifying additional genes and functions. We also discovered a subset of risk variants interacting with multiple promoters and the expression levels of such genes were correlated. The presented resource enables functional studies of cardiovascular disease providing novel approaches for its diagnosis and treatment.

## Introduction

Cardiovascular disease (CVD) labels medical problems of the circulatory system (heart, blood vessels and arteries) often due to build-up of fatty cell debris (plaques) deposited inside the blood vessels. It is the leading cause of disability and death globally ^1^. Atherosclerosis, the main underlying mechanism leading to the acute events of CVD, is characterized by a lipid driven chronic inflammation of the arterial intima, a process that includes all major cells in the vascular wall, i.e. endothelial, smooth muscle and inflammatory cells. The acute complication of atherosclerosis such as myocardial infarction and stroke is due to rupture of the fibrous cap with subsequent thrombus formation that totally or partially occludes the vessel and thereby stops the nutrient-rich blood flow. Traditional risk assessment methods based on age, gender, smoking, diabetes, hypertension and dyslipidemia tracks the disease incidence well but underestimates its occurrence since almost half the population classified as low or intermediate risk end up developing cardiovascular disease ^2-4^ as these methods fail to inform on the underlying pathological processes that may have been going on for years ^5^. In addition, ethnic differences in cholesterol and blood lipid levels complicate the assessment of the individual risk of disease ^6^. Heritability for CVD hovers around 0.5, therefore genetic risk contributors at play can be utilized in its early diagnosis and treatment ^7^.

Genome-wide association studies (GWAS) have emerged as an important tool in the search for disease causing genomic variants ^8^. Coronary artery disease-specific and other atherosclerosis related indications have been addressed by large GWAS meta-analyses enabled by consortia such as the CARDIoGRAMplusC4D ^9^ and the MEGASTROKE consortium ^10^. At its current state, just over 300 independent variants explain 21% of coronary artery disease heritability ^11^. According to GWAS, a locus on chromosome 9p21 has the strongest association signal ^12,13^. Although it is established that the risk allele is associated with formation and progression of plaques but not with their rupture ^14,15^, the mechanistic understanding of the conferred risk by these loci remains elusive ^16-18^. Pathways such as cholesterol and triglyceride metabolism, blood pressure, inflammation, vascular proliferation and remodeling, nitric oxide signaling, vascular tone, extracellular matrix integrity and axon guidance and signaling are also enriched for target genes of GWAS variants ^17-20^.

GWAS studies do not in themselves provide functional insight for the large subset of hits that are non-coding ^21,22^: only one-third of the time a variant affects the expression level of its nearest gene, highlighting the limitations of the nearest-gene assignment approach ^23,24^. The target gene mappings can be refined using various layers of genome annotation information as well as gene expression profiles. To alleviate the problem of complex linkage structures between variants, vast amounts of public datasets of epigenetic marks and transcription factor binding profiles employed to help prioritize the causal/functional variant ^25-27^. Expression quantitative trait loci (eQTL) analyses based on gene expression and genotype datasets are also used to locate potentially functional variants that are in linkage disequilibrium (LD) with top association variants ^28-30^.

Pathway or gene-set based approaches using canonical pathways and gene ontology (GO) terms goes beyond single variant-based analyses and investigate the combined effect of multiple disease/trait variants on biological functions in terms of the perturbations on pathways or cellular processes ^31-35^. Such pathway based analyses revealed the functional GWAS variants in cases such as Crohn’s disease ^36^, multiple sclerosis ^37^, schizophrenia ^38^ and breast cancer ^39^. Functional gene sets built using co-expression and protein-protein interaction datasets are also used successfully to interpret the GWAS variants ^40-42^.

Studies of the chromatin interaction landscape were revolutionized by the invention of chromosome conformation capture coupled with next-generation sequencing (Hi-C) ^9^. Combining Hi-C with sequence capture (HiCap), the improvement in resolution required for study of individual promoter-enhancer interactions can be obtained ^43,44^. In this study, we employed HiCap on three cell types relevant for cardiovascular disease to discover novel biological processes and pathways related to onset and pathology of the disease. We utilized chromatin contacts of promoters to GWAS variants or those that are in linkage disequilibrium (LD) to assign potential target genes.

## Results

Using high-resolution chromatin interactions, we mapped genomic interaction of promoters and variants associated with traits and conditions related to cardiovascular disease. We employed three cell types for the investigation: human aortic endothelial cells (AEC), human aortic smooth muscle cells (ASMC) and macrophage-THP-1 cells (mTHP-1). For AEC, we obtained two technical replicates from two individuals; the technical replicates were pooled, and the two individuals were held separate and constituted biological replicates. For ASMC and mTHP-1 cells, two technical replicates were obtained. We employed HiCap using a probe set targeting 21,479 promoters and 3,950 variants (suppl. table 1). There were 199, 300 and 176 million read pairs uniquely mapped to probes in AEC, ASMC and mTHP-1 experiments, respectively (suppl. table 2). We made a distinction between interactions between promoters (promoter-promoter or P-P) and those between promoters and elsewhere in the genome (promoter-distal or P-D). For the sake of clarity, we also defined interactions between disease/trait associated variants and promoters (GWAS-Promoter or G-P) (figure 1a).

**Figure 1.**
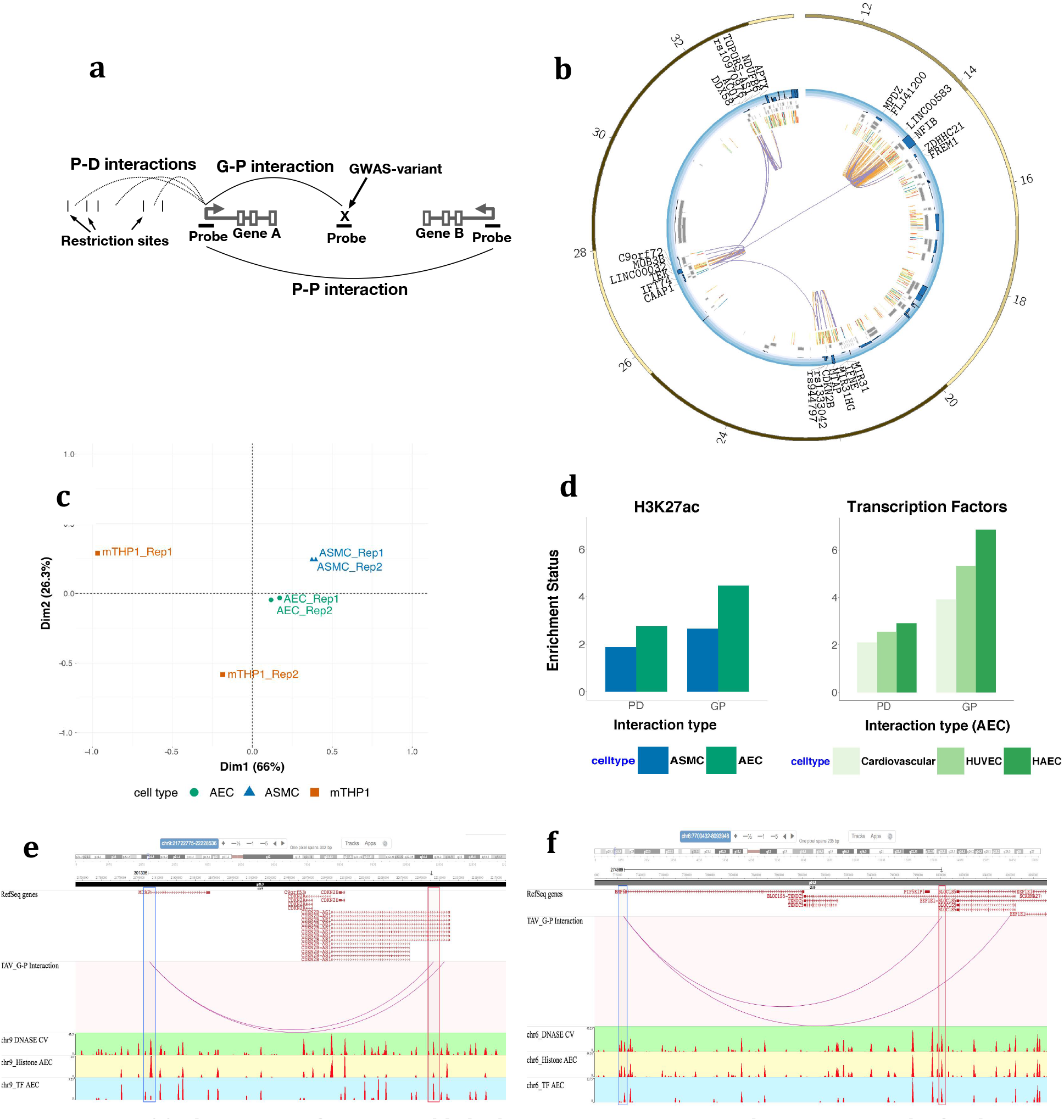
(a) Three types of HiCap-established interactions; P-D, P-P and G-P, respectively; for the Gene A promoter. In P-D is analyzed probed promoters interaction with distal element (DEs); the latter separated by restriction sites. In P-P and G-P both ends of the interaction are probed. (b) The largest connected subgraph in chr9 in AEC cells. There are 22 promoters, three GWAS SNPs and 271 DE (only those overlapping with H3K27ac marks are visualized). The blue track represents gene expression levels, gray boxes represents transcripts, innermost layers represent H3K27Ac marks (two replicates). Purple arcs represent G-P or P-P, orange arcs represent P-D interactions. (e,f) Interactions between, respectively (c) Principal component analysis (PCA) of P-D interaction of genes that are not expressed in all cell types. The supporting pairs (SP) read counts were normalized with respect to distal element length and PCA was performed. (d) Overlap enrichment relative to a segment-length and distance-from-promoter controlled random set. The AEC P-D and G-P datasets were overlapped with general Cardiovascular, HUVEC and HAEC transcription factor marker data from ChipAtlas. (e) MTAP-rs944797 and (f) BMP-rs9328448 as visualized with the WashU browser. Overlap with CV-DNAse, HAEC-Histone and HAEC-TF markers from ChipAtlas are shown in the lower panes.

We called interactions using HiCapTools ^15^, and p-value cutoffs deployed yielded interaction sets of sizes 69,753 (AEC P-D), 38,759 (ASMC P-D), 20,699 (mTHP-1 P-D), and 5,671 (AEC P-P), 4,293 (ASMC P-P) and 8,921 (mTHP-1 P-P), respectively (suppl. table 3, suppl. methods). Importantly, we were able to detect many long-range (>500Kb) interactions across the three cell-types (suppl. figure 1a). Equally important, the distal elements (DEs) as well as the interacting promoters were short; average length being 749 and 776 bases, respectively. In total, the interaction datasets covered around 3.3% of the genome. Most promoters (65%) were found to interact with less than five distal regions, whereas the interactome of extreme hub-promoters contain several hundred DEs (suppl. fig. 1b-d). We identified several interconnected units of promoters and enhancers, figure 1b displays the largest connected subsection of chromosome 9 (i.e. giant component). Interestingly, two CVD associated GWAS SNPs in chr9p21 region (rs1333042 and rs944797) were part of this network.

Utilizing principal component analysis, we show that the interaction profiles of individual cell types are specific and can separate individual cell types independent of gene expression information (figure 1c, suppl. table 4, suppl. figure 3).

### Promoter-interacting distal elements were enriched for functional elements

Promoter-interacting DEs were previously shown to be highly enriched for enhancer marking features ^12,14,15^. To confirm that is also the case for this study, we overlapped DEs with H3K27ac enriched regions obtained through ChIP-seq in the same cells, as well as relevant DNaseI, histone modification (H3K27ac and H3K4me1) and transcription factor binding datasets from the ChipAtlas (chip-atlas.org). Our interactor sets were indeed enriched relative to size- and genomic context controlled random sets and the enrichment was stronger for the better matching cell types (figure 1d, suppl. figure 4a-d). Interestingly, enrichment levels for the G-P set was much higher in AEC and ASMC but not in mTHP-1 cells (figure 1d, suppl. figure 4a-d). Further, promoters interacting with DEs carrying H3K27ac marks were expressed at higher levels as expected (suppl. fig 4e).

Figure 1e and 1f shows two examples of promoter interactions (MTAP-rs944797 and BMP-rs9328448) where the interactor overlaps with both CVD GWAS variants and enhancer marks.

### Promoter interacting GWAS-variants were often contained within regulatory elements

We next turned our attention to variants associated with cardiovascular disease phenotypes according to GWA studies and asked if any variants or in LD with those are contained within DEs found in this study. First, we took all single nucleotide variants associated with “cardiovascular disease” resulting in 3,814 SNPs (minimum p value for association is 10^−6^), which we called CVD_GWAS ^45^ (suppl. table 5). As previously shown, a substantial portion of GWAS variants are themselves located within potential regulatory regions ^46^. We therefore targeted a subset (723, 19%) of these variants using probes to increase the probability of obtaining a signal without the need for deep sequencing (suppl. table 5). Of those 723 targeted GWAS variants, 314 (43%) interacted with at least one promoter in at least one cell type, constituting the G-P dataset. We assigned 512 target genes to 314 variants (42% interacting with only one promoter, 20% with two promoters). To rule out the possibility that G-P hits occur merely as result of the probing of variants we investigated the corresponding P-D set, i.e. we studied the same promoter and its interaction with non-probed distal regions. There should be vastly more P-D hits close to the variant site, than to a site on the same distance from the promoter but at the other side of it (hence keeping the distance from the probed feature the same). This is indeed the case as shown in figure 2a.

**Figure 2:**
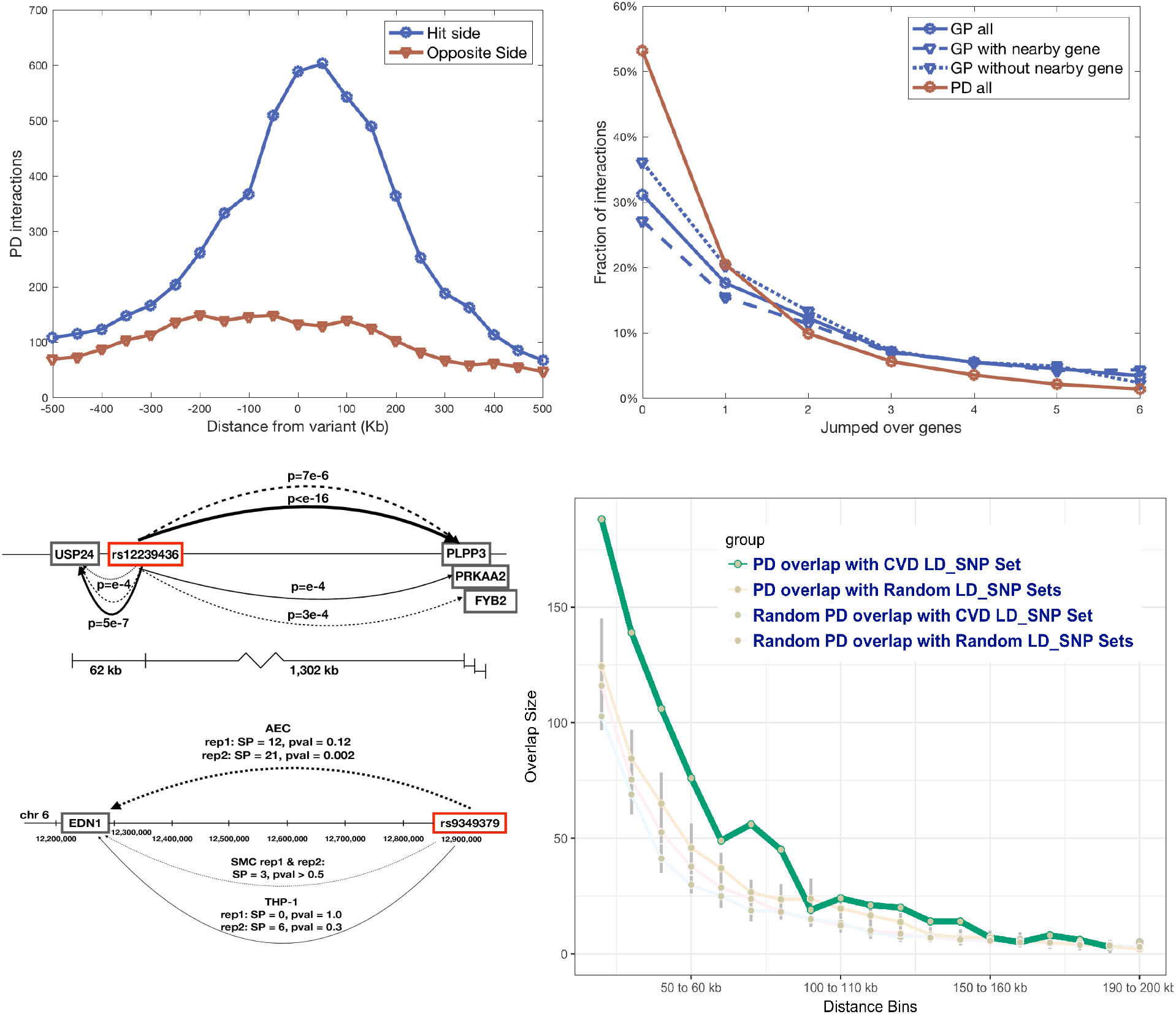
(a) The AEC P-D set was searched for interactions between the promoter and distal elements close to a variant interacting with the same promoter (blue curve). The comparison was made relative to a site at the same distance from the promoter it but located at the other side of it (red curve). The latter is nearly a horizontal line as expected whereas the blue curve strongly deviates from that at distances not too far from the variant site. (b) Variants in the AEC, ASMC and mTHP-1 merged G-P set and their interaction preferences with genes at distance 0 (no gene-jumping), 1 (nearest gene is jumped-over) etc. Distal regions in the corresponding P-D set are shown for reference. The G-P interaction set is further split in equally sized halves depending the variants distance to its nearest gene (G-P without- and with a nearby gene, respectively). (c) The CAD-related variant rs12239436 (red box) interacts with 62 kb distant gene USP24 (grey box) in AEC (dotted line), ASMC (dashed) and mTHP-1 (solid). Strengths of interactions are represented with the p-value recorded and indicated by arrow thickness. Our result set further include very strong interactions with the 1,3Mb distant previously CAD associated gene PLPP3 (also known as PPAP2B). Less significant interactions with nearby FYB2 (ASMC) and PRKAA2 (mTHP-1) are potentially bystanders. (d) Earlier reported interaction between variant rs934937 and the EDN1 gene is, to a varying degree, present in both AEC patients. Not so in less relevant tissues ASMC and mTHP-1. (e) Comparison of overlaps between P-D dataset (AEC) vs 100 matched random datasets and CVD_GWAS and matched SNP datasets (see methods) shows that there is an enrichment for variants in LD with CVD_GWAS found in P-D dataset when the genomic distance between SNP and its LD proxy is <= 80 kb. No such enrichment was seen for random P-D datasets vs real or random SNP sets.

A large fraction of G-P interactions spanned distances above 500 kb. Consequently, many of them preferentially interact with non-closest genes thus ‘jumped over’ by the interaction loop formed. Indeed, only less than a third interact with adjacent genes as shown in figure 2b. For the P-D interactions the fraction is larger, about 55%. Not unexpectedly the gene jumping among G-P interactions is even more frequent when the closest gene is near in distance.

Whole groups of genes frequently interact with the same GWAS variant. Promoters of genes USP24, PLPP3, PRKAA2 and FYB2 thus share a putative enhancer containing variant rs12239436. The rs12239436-PLPP3 interaction is particularly interesting due to its large 1.3Mb distance as well as the fact that PLPP3 was already identified as a CAD disease risk gene ^47^ (figure 2c).

The GWAS variant rs9349379, associated with five vascular diseases, was recently shown to regulate expression of the Endothelin-1 gene ^48^. We see this interaction in one of the two AEC investigated individuals. In cell lines ASMC and mTHP-1 the interaction is weak (figure 2d).

### Discovery of target genes of GWAS variants using shared haplotype and interaction information

To detect further interactions of CVD_GWAS variants with promoters, we looked at the fraction of DEs containing such variants. Of the 3,814 associated variants in the CVD_GWAS, there were 150 (3.9%) variants within unique DEs. One complication of GWA studies is that the association signal many GWAS variants possess are due to their sharing of haplotype with the functional variants. If the functional variant can indeed modulate a distal promoter via looping, it should also be possible to locate it in our DE datasets. We therefore looked at the fraction of DEs that contain variants that are in LD with those in CVD_GWAS. Due to the sheer number of SNPs in LD, we devised a double randomization scheme to assign statistical significance to the observed overlap between LD SNPs and DE datasets using both size- and context-matched random interaction datasets and random SNP datasets matched with respect to allele frequencies and surrounding LD structure of the real set (suppl. methods). Figure 2e shows that the DE dataset is enriched for SNPs in LD with CVD_GWAS that are within 80 kb and 20 kb in AEC and ASMC respectively, beyond which no enrichment can be seen (suppl. figure 5a). mTHP-1 cells showed limited enrichment (suppl. figure 5b). There were 331 SNPs in LD with variants in CVD_GWAS located within DE fragments (P-D_LD). DEs carrying either the GWAS variant themselves or those in LD showed higher enrichments for open chromatin, transcription factor binding sites and enhancer marks compared to the entire DE set, supporting their potential for expression modulation (figure 3a).

**Figure 3.**
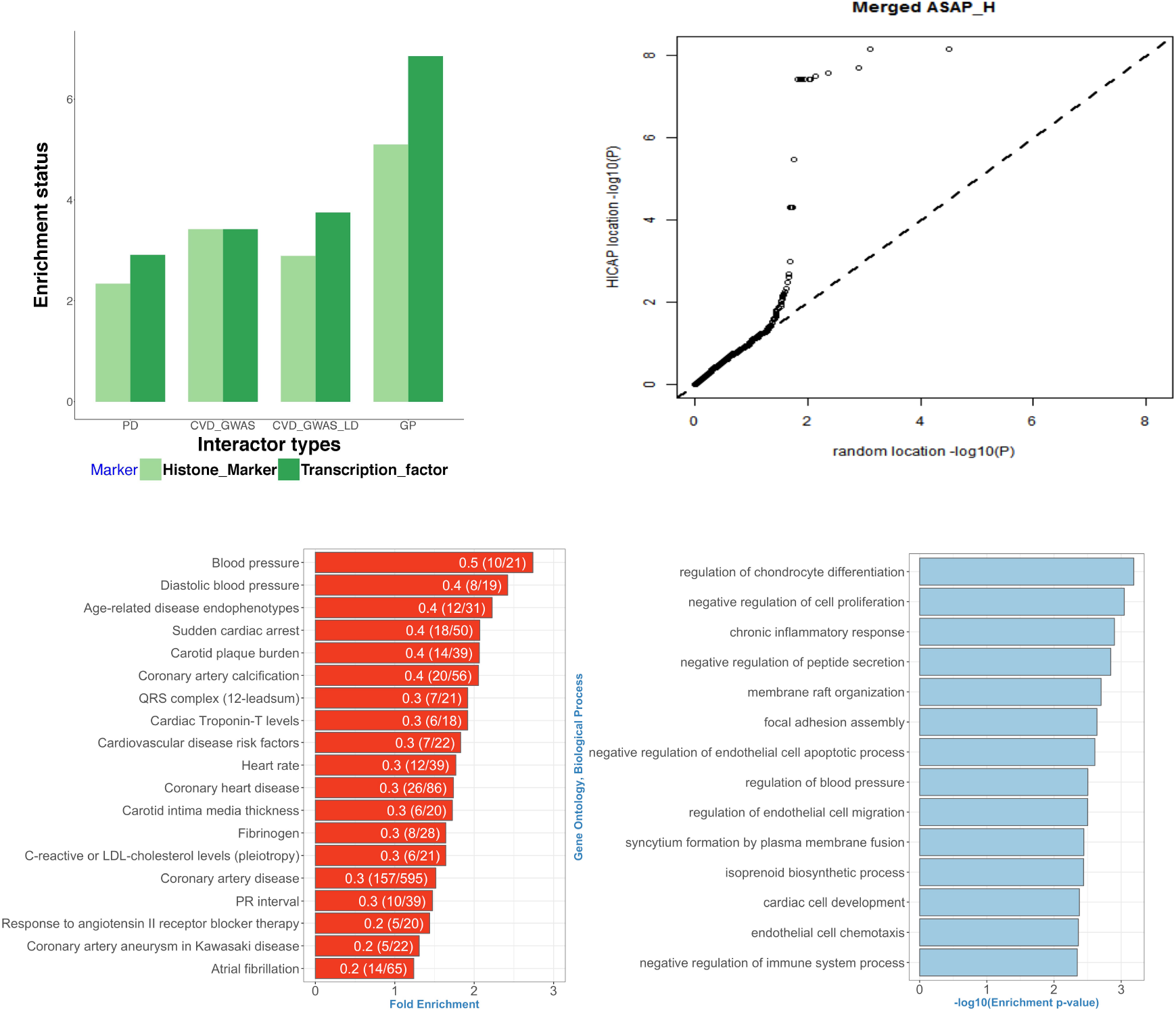
(a) Overall enrichment of the P-D (AEC) datasets to the functionally annotated transcription factor and histone markers from Human Aortic Endothelial cell (HAEC) chip-atlas database. The enrichment status was assigned with respect to segment-length and distance-from-promoter controlled random set datasets. (b) eQTLs contained in the merged CVD G-P, P-D and P-D_LD datasets plotted versus a size and distance-corrected random set. Deviation from diagonal is present among approximately 30% of the data. (c) GWAS traits that are overrepresented in the interaction datasets. Only traits containing at least 14 variants were taken forward. Fold enrichment is calculated by dividing the actual number of trait variants in the interaction dataset to that of expected (fraction of trait variants in the full trait set). The bar labels denote the fraction of variants found in the interaction datasets. (d) GO Term enrichment analysis of genes interacting with variants or those in LD with CVD_GWAS set using TopGO package. GO terms enriched using only nearest genes to the variants are not reported. Terms with less than five genes and enrichment score less than 0.005 were not included.

The eQTL technique was deployed to examine the expression modulation capacity of variants contained in DEs. The G-P set was extended with P-D hits and likewise selected hits in linkage disequilibrium with those (P-D_LD). The comparison was performed relative to the afore mentioned size- and distance controlled random set and yielded the Q-Q plot presented in figure 3b (merged set) and suppl. figure 6a-c (G-P, P-D and P-D_LD separate). The deviation from the diagonal is striking and concerns not only extreme cases but large fractions of the entire sets.

Only 27% of the DEs containing all these variants interacted with the closest gene and the average interaction distance is 358 kb (suppl. table 6). In terms of trait categorization, variants for 145 of the 247 traits (0.59) related to CVD in EBI GWAS catalogue was found, figure 3c lists overrepresented traits in our dataset.

### Gene enrichment analysis of target genes of GWAS variants for discovery of disease associated cellular processes

We next asked if genes interacting with fragments carrying disease associated variants are enriched for particular functions or pathways. To discover cell context dependent signal, we performed a gene set enrichment analysis using genes interacting with GWAS variants themselves and those in LD for each cell type. We only included LD SNPs up to 80 kb and 20 kb to the proxy SNP in AEC and ASMC cells and no LD SNPs were included in mTHP1 cells. In order to assess the success of discovering novel processes or functions, we input the DEs containing these variants to GREAT software package to retrieve the gene sets independent of interaction information to perform the same enrichment analysis for comparison ^49^ (suppl. methods). We performed enrichment analysis separately for each cell and also combined to assess the specific contribution of each cell type. Comparison of the enriched terms by interacting or closest gene information (GREAT package) using a GO term semantic similarity measure revealed little overlap in between (suppl. methods, suppl. table 7) ^50^. Figure 3d shows enriched biological processes when target genes from all cell types are merged. Only one term (“positive regulation of transcription, DNA-templated”) were similar to those found using closest gene information (suppl. table 7). We located novel genes associated with known CVD complications including “response to lipopolysaccharide” (GO:0032496, in AEC and mTHP1), “phosphatidylinositol-3-kinase signaling”(GO:0014065, in AEC) and “SMAD protein signal transduction” (GO:0060395, in AEC and ASMC). Moreover, we discovered genes and functions not previously associated with CVD onset and or progress such as “cilium assembly” (GO:0060271, in AEC, 14 genes).

Ectopic deposition of calcium in arterial vessel walls leading to vascular calcification is a main feature of atherosclerosis ^51^ and similar to the ossification process ^52^. Concordantly, terms such as “endochondral ossification”, “regulation of chondrocyte differentiation”, “regulation of osteoblast differentiation”, “positive regulation of ossification” were among the enriched functions. Eleven genes (BMP6, SMAD3, JAG1, PDLIM7, SLC8A1, DLX5, TEK, HOXA2, EFEMP1, RARB and IL6) responsible for the above enrichments and only four (BMP6, TEK, SMAD3 and JAG1) interacted with lead SNPs, while the rest interacted with variants in LD with lead SNPs.

Fourteen target genes were involved in cilium assembly, including IFT74, a component of endothelial intraflagellar transport ^19^, which interacts with a CVD associated variant in the chr9p21 region (rs944797). It has been shown that endothelial cells can sense and respond to shear stress levels using their cilia ^19,20^ and endothelial cilia were shown to deflect in response to blood flow rates. The deflection angle is regulated by calcium levels ^21^. Moreover, endothelial cilia inhibit onset of atherosclerosis in mouse models ^22^.

We identified several genes involved in leukocyte adhesion and vascular inflammation, key processes of atherosclerotic development. Examples of target genes include Cadherin 13 (CDH13 interacting with rs8055236) which has previously been shown to protect against atherosclerosis in experimental models 52, AMP-activated protein kinase (PRKAA2 interacting with rs12239436) whose activity inhibits cell migration via phosphorylation of Pdlim5 ^53^ and BACH1 (interacting with rs2832227), a transcriptional regulator which has been shown to be involved in atherosclerosis development in apolipoprotein E deficient mice ^54^.

Other examples of plausible candidate genes for inflammatory cardiovascular disease include CD86 (interacting with rs13083990), a receptor involved in the costimulatory signal essential for T-lymphocyte proliferation and interleukin-2 production ^55^ and AKIRIN2 (interacting with rs6900057) a gene that has been shown to stimulate a pro-inflammatory gene in macrophages during innate immune responses ^55^.

### Expression levels of interacting promoters sharing enhancers are correlated

Sometimes variants are contained within enhancers controlling multiple genes, suggesting such gene sets to be group- and pairwise co-expressed. Using data from the ASAP-Heart study we were able to test 75 AEC gene pairs sharing enhancers and could conclude that 34 (45.3%) are co-expressed at p-value level 10^−3^; 18 of 75 (24.0%) also at p-value level 10^−10^. Excluding a large cluster of genes all interacted upon by the same variant rs13083990, these percentages rises to 64.1% (p < 10^−3^) and 38.4% (p < 10^−10^), respectively.

In figure 4a this is exemplified with co-expression plots for the MTAP and IFT74 genes, both interacted upon by the chr9p21 locus variant rs944797 in AEC. The genomic distance between the two gene promoters is in excess of 5 million bases. Even more extreme is the co-expression of genes SMARCAD1 and BMP2K both located on chromosome 4 more than 15 Mb apart in AEC (figure 4b). Both genes also interact a GWAS variant associated with diastolic blood pressure (rs16998073, p value = 10^−21^) ^56^, which itself interacts two other genes (LINC01094 and PAQR3) and 27 DEs (11 overlapping with H3K27ac marks) (figure 4c). Most of these interactions were specific to AEC (figure 4d). LINC01094 is a non-coding RNA and its expression levels are correlated with serum albumin (p value =10^−9^) ^57^ and co-expressed with BMP2K (p value = 1.26 * 10^−93^). Serum albumin levels are positively correlated with blood pressure ^58^. Moreover, PAQR3 levels modulates leptin signaling in mouse models ^59^ and leptin is found to mediate the increase in blood pressure associated with obesity ^60^.

**Figure 4.**
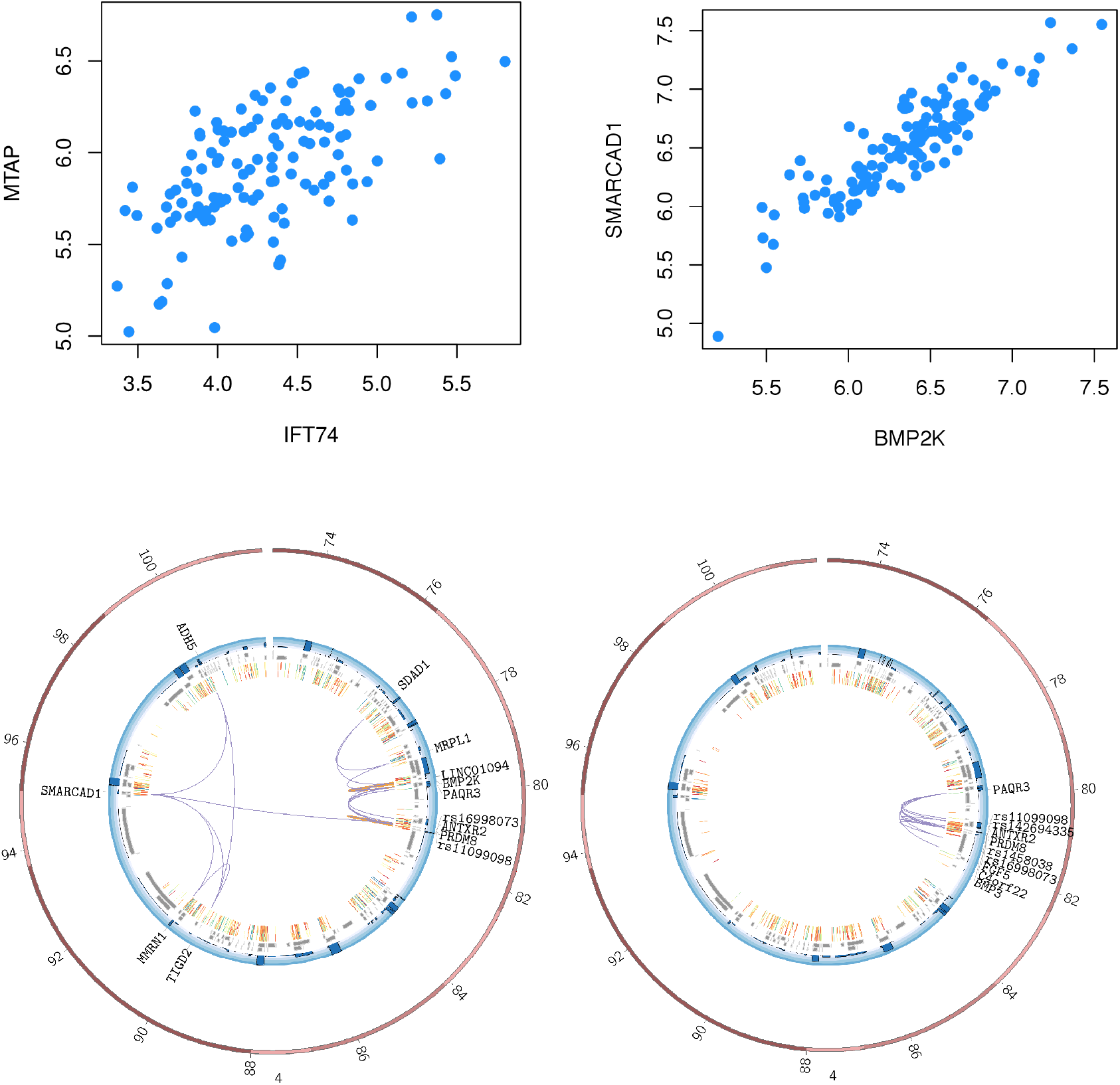
Expression correlation between (a) genes MTAP and IFT74 (p-value 4.2*10^−16^) and (b) genes SMARCAD1 and BMP2K (p-value 2.1*10^−45^) using aortic intima media expression from 131 individuals. c,d) Circos plot representation of interactions between rs16998073 and rest of the genome in c) AEC and d) ASMC for comparison. The plot spans chr4:73000000-103000000. The blue track represents gene expression levels, gray boxes represents transcripts, innermost layers represent H3K27Ac marks in AEC cells (two replicates). Purple arcs represent G-P or P-P and orange arcs represent P-D interactions.

## Conclusions

Our aim in this study was to evaluate the contribution of high-resolution promoter-anchored regulatory interaction maps to locate the target genes of noncoding GWAS variants associated with cardiovascular disease. Although many GWAS variants are merely tags for the functional SNP within the same haplotype, some may still be or be very close to the “functional” variant as suggested by their enrichment for enhancer marks and re-sequencing studies ^46^. Here we show that targeting GWAS variants in capture Hi-C experiments can be a useful strategy in conquest for target gene associations due to lesser need for sequencing. We also uncover several enhancers regulating multiple genes and a strong correlation signal between such sharing enhancers, implying the underlying complexity of regulatory networks. A recent study showed the implicit wiring of enhancer redundancy in regulatory networks ^61^, where the system can tolerate loss of enhancers by connecting promoters to multiple deplorable enhancers. However, the case when multiple genes connected to the same enhancer could negatively affect the resilience of the network in the case of enhancer malfunction, potentially disturbing the co-regulation of multiple genes.

We tackle the difficulty of locating the “functional” variant using LD information. When a GWAS SNP is associated with a trait, essentially any other variant on the same haplotype could be responsible for the association. However, due to sheer number of variants in LD, it is not straightforward to locate the “functional” one. The resolution in this study was around 750 bases, which allowed us to locate DEs containing variants in LD with CVD GWAS variants. Extending the genomic window around the DEs to 200 kb, we found that, it is possible to discriminate between functional and tagged variants using a double randomization procedure. DEs containing variants in LD with CVD GWAS variants showed better enrichments for histone enhancer marks and TF binding sites and are more likely to contain eQTLs.

We confirm that it is only one-third of the time the enhancer is connected to the promoters of its nearest gene. We take on the challenge of assigning the correct genes to GWAS variants using promoter- and variant-anchored regulatory maps produced in three cell types. Indeed, we discover multiple biological processes and cellular structures that are associated with cardiovascular disease pathology not by genomic but by functional studies. We were able to suggest variants that could be responsible for the perturbations of such processes or structures. Here it is important to note that by only mining the variants associated with cardiovascular disease trait, we will be able to discover the genes that are perturbed in a given pathway, process or structure. Discovery of the full network of genes within such processes or structure is beyond the scope of this study.

In summary, we provide high-resolution promoter-anchored regulatory networks of three cell types and list novel genes, processes and cellular structures relevant for cardiovascular disease pathologies. We hope that the data and the methodologies in this study will aid us in our mission to further understand the contribution of non-coding genomic variation to complex disease biology.

## Methods

Please see supplementary material for detailed method descriptions.

